# SMAS: Structural MRI-Based AD Score using Bayesian VAE

**DOI:** 10.1101/2024.11.07.622411

**Authors:** A. Nemali, J. Bernal, R. Yakupov, D. Singh, M. Dyrba, E.I. Incesoy, S. Mukherjee, O. Peters, J. Hellmann-Regen, E. Ersözlü, L. Preis, J. Priller, E. Spruth, S. Altenstein, A. Lohse, A. Schneider, K. Fliessbach, O. Kimmich, J. Wiltfang, N. Hansen, B. Schott, A. Rostamzadeh, W. Glanz, M. Butryn, K. Buerger, D. Janowitz, M. Ewers, R. Perneczky, B. Rauchmann, S. Teipel, I. Kilimann, D. Goerss, C. Laske, S. Sodenkamp, A. Spottke, M. Coenjaerts, F. Brosseron, F. Lüsebrink, P. Dechent, K. Scheffler, S. Hetzer, L. Kleineidam, M. Stark, F. Jessen, E. Duzel, G. Ziegler

## Abstract

This study introduces the Structural MRI-based Alzheimer’s Disease Score (SMAS), a novel index intended to quantify Alzheimer’s Disease (AD)-related morphometric patterns using a deep learning Bayesian-supervised Variational Autoencoder (Bayesian-SVAE). SMAS index was constructed using baseline structural MRI data from the DELCODE study and evaluated longitudinally in two independent cohorts: DEL-CODE (n=415) and ADNI (n=190). Our findings indicate that SMAS has strong associations with cognitive performance (DELCODE: r=-0.83; ADNI: r=-0.62), age (DEL-CODE: r=0.50; ADNI: r=0.28), hippocampal volume (DEL-CODE: r=-0.44; ADNI: r=-0.66), and total grey matter volume (DELCODE: r=-0.42; ADNI: r=-0.47), suggesting its potential as a biomarker for AD-related brain atrophy. Moreover, our longitudinal studies suggest that SMAS may be useful for early identification and tracking of AD. The model demonstrated significant predictive accuracy in distinguishing cognitively healthy individuals from those with AD (DELCODE: AUC=0.971 at baseline, 0.833 at 36 months; ADNI: AUC=0.817 at baseline, improving to 0.903 at 24 months). Notably, over a 36-month period, SMAS index outperformed existing measures such as SPARE-AD and hippocampal volume. Relevance map analysis revealed significant morphological changes in key AD-related brain regions—including the hippocampus, posterior cingulate cortex, precuneus, and lateral parietal cortex—highlighting that SMAS is a sensitive and interpretable biomarker of brain atrophy, suitable for early AD detection and longitudinal monitoring of disease progression.

## I. Introduction

Alzheimer’s Disease (AD) represents a major global health challenge, largely due to its increasing prevalence among the aging population and its substantial socioeconomic implications. With demographic changes leading to increased prevalence, AD and other age-related dementias are projected to significantly strain healthcare systems and economies worldwide [1]–[4]. Therefore, it’s crucial to prioritise the development of accurate and early diagnostic tools for AD. A promising approach is the development of a risk assessment score to quantify disease stage in its early phases. Such scores could facilitate early detection, enabling timely interventions at stages when treatments are most effective. In addition, these scores could be extremely valuable in identifying patients and stratifying them for clinical trials, ensuring that research is effectively targeted and adapted to the specific stages and needs of patients, leading to more efficient and personalized care for those affected by AD [5]–[7].

The National Institute on Aging and Alzheimer’s Association (NIA-AA) Research Framework defines AD by three key biomarkers: amyloid (A), tau (T), and neurodegeneration (N). These markers reflect the pathological processes that occur in the brain during AD progression [8]. Among these biomarkers, neurodegeneration can be detected using structural Magnetic Resonance Imaging (sMRI), a non-invasive tool that offers greater accessibility compared to Positron Emission Tomography (PET) and Cerebrospinal Fluid (CSF) analysis, which are typically used for detecting amyloid and tau proteins [9], [10].

A critical challenge is developing a risk assessment score that characterizes AD-related brain atrophy patterns using sMRI. Traditional approaches to quantify brain atrophy related to AD often rely on regions affected such as hippocampal atrophy (a prominent biomarker of AD) and entorhinal cortex measurements [53]–[55]. However, they might lack specificity for AD and struggle with sensitivity and specificity at the mild cognitive impairment (MCI) stage, which can also manifest in non-AD dementias [56]–[58]. Therefore, solely relying on these regions for a risk assessment score may overlook the comprehensive spatio-temporal patterns of brain atrophy associated with AD.

A data-driven machine learning approach involves the extraction of morphometric features from sMRI scans, such as tissue segments (gray matter (GM), white matter (WM) & CSF), can reveal patterns of brain changes associated with aging and AD, potentially predicting disease progression and severity [13], [14]. However, the high dimensionality and heterogeneity of these imaging features present significant challenges in the analysis, often limiting the effectiveness (in terms of performance) of traditional approaches [15], [16]. One promising approach involves identifying low-dimensional latent representations of large, high-dimensional datasets. Numerous studies have employed multivariate approaches such as support vector machine (SVM) classifier to estimate an index of AD anatomical risk called the spatial pattern of brain abnormality for recognition of early AD (SPARE-AD) [16], partial least squares (PLS), canonical correlation analysis (CCA), and their sparse variants (sPLS and sCCA). These techniques have been applied to associate cognitive scores or symptoms with imaging data [17], [18] and for multimodal analysis [19], [20]. However, these linear models might fall short in capturing the intricate interplay and non-linear dynamics that characterise AD progression [21]–[23].

Advancements in multimodal deep learning have shifted the focus towards models that can learn these non-linear relationships and extract meaningful features from complex data [24]–[26]. Variational Autoencoders (VAEs) [27], [28], a class of generative models, have shown promise in learning highlevel probabilistic latent embeddings from data, facilitating the identification of common patterns in aging and AD progression [29], [30]. However, the complexity and heterogeneity inherent in sMRI scans used in AD research pose challenges for VAEs. Issues such as posterior collapse, i.e. when the learned latent variables become too similar to the prior, leading to a failure in capturing meaningful variability such as brain atrophy, and lack of supervision can impede the VAE’s effectiveness in extracting meaningful latent representations from sMRI data [31].

To overcome these challenges, we here focus on a supervised VAE [28]. Utilizing this approach we introduce a supplementary variable (or prediction node) that accounts for the conditional relationships between morphometric features and a cognitive score that is related to AD disease progression. By integrating cognitive performance differences directly into the model’s architecture, the VAE learns a more relevant representations of the imaging data and potentially improving specificity in identifying AD-related change patterns that are conditioned on specific cognitive states. Therefore, this approach could be useful for improved quantification of individual disease progression or staging in AD.

In medical image analysis, the reliability of ML models is critical for supporting quantification of relevant uncertainties and transparency. Our approach focuses on three key aspects: uncertainty quantification, explainability, and generalizability, to improve model robustness and interpretability. To address uncertainty arising from inter-subject variability, scanner heterogeneity, and model complexity [34], [36], we implement a Bayesian framework within the supervised VAE, treating model parameters as random variables with prior distributions, thereby enhancing predictive accuracy and transparency. For explainability, we utilize gradient activation maps to identify the image regions most influential to the model’s predictions, offering insight into the decision-making process employed by the model, thereby facilitating clinicians’ understanding and trust in the rationale behind its predictions. Generalizability is achieved through validation on independent observations, testing the model’s performance across datasets (including variation scanner resolution), thus supporting its broader clinical applicability.

More specifically, in this study we propose a Structural MRI-based Alzheimer’s Disease Score (SMAS) using a novel Bayesian supervised Variational Autoencoder model (Bayesian-sVAE). The SMAS was derived from sMRI images and conditioned on cognitive scores, with the aim of capturing unique patterns of brain atrophy associated with AD-related cognitive impairments. The study utilized longitudinal sMRI data from two large multi-center cohorts: the DELCODE (DZNE Longitudinal Cognitive Impairment and Dementia) study, which is part of the German Center for Neurodegenerative Diseases (DZNE) [37], and the Alzheimer’s Disease Neuroimaging Initiative (ADNI) study (adni.loni.usc.edu). The analysis follows two main stages: 1) Training the Bayesian-sVAE model on baseline DELCODE data, which included subjects from various diagnostic groups: cognitively normal (CN), subjective cognitive decline (SCD), MCI, AD, and AD-relatives (ADR), allowing the estimation of SMAS index; and 2) Validating the trained model using the two datasets: the unseen DELCODE follow-up data and the independent ADNI dataset. Finally, the estimated SMAS index underwent rigorous testing through several analyses, including A) correlating the SMAS index with various clinical assessments; B) tracking the longitudinal progression of the SMAS index; (C) evaluating the SMAS index in subjects who remained unimpaired or became impaired during the study; and D) comparing the SMAS index with the well-established SPARE-AD [16] index and evaluating their performance in classifying CN subjects versus those with MCI or AD.

## II. Materials & Methods

### A. Bayesian Supervised Variational Autoencoder

Variational Autoencoders (VAEs) provide a powerful probabilistic framework for analyzing sMRI features by combining deep learning with variational inference. VAEs are particularly suitable for capturing the non-linear dynamics associated with AD [45]. In this context, let *x* represent MRI-based (potentially preprocessed) structural images (or input features) that capture neuroanatomical changes, and let *z* denote a latent (or hidden) variable. The VAE is designed to learn the joint distribution *p*(*x, z*), which can be decomposed as *p*(*x, z*) = *p*(*z*)*p*(*x*| *z*), where *p*(*z*) is the prior distribution over latent variables and *p*(*x*| *z*) is the likelihood of the sMRI imaging features given the latent representation. The latent variable *z* is assumed to capture relevant aspects of brain atrophy patterns present in the sMRI images.

In this study, we propose a Bayesian Supervised Variational Autoencoder (Bayesian-sVAE) generative model to learn latent representations of brain structure from MRI features while simultaneously predicting a dependent variable, such as cognitive scores or clinical diagnoses. The Bayesian sVAE extends the traditional sVAE framework by treating the model parameters as random variables and employing variational inference for Bayesian model optimization.

The Bayesian sVAE model consists of three main components: an encoder **q**_*ϕ*_(**z**|**x**), a decoder **p**_*θ*_(**x**|**z**), and a regressor **q**_*ψ*_(**y**|**z**), where **y** represents the dependent variable. The encoder network transforms the input MRI samples **x** = [*x*_1_, *x*_2_, …, *x*_*n*_] and dependent variables **y** = [*y*_1_, *y*_2_, …, *y*_*n*_] into a lower-dimensional latent space, where each *x*_*i*_ rep-resents MRI features of the *i*^*th*^ subject. We assume that each MRI feature *x*_*i*_ is linked to a latent representation *z*_*i*_, and **z** = [*z*_1_, *z*_2_, …, *z*_*n*_] is a latent vector influenced by the dependent variable **y**.

The **encoder network q**_*ϕ*_(**z**|**x**) comprises a 3D convolutional block, several Residual Network (ResNet) blocks, and a fully connected layer. The ResNet blocks include two 3D convolutional layers with batch normalization and ReLU activation, along with a skip connection. As the network deepens, the number of filters in these blocks increases progressively (for more details see [48]). The fully connected layer produces a latent vector *z*_*i*_ of *m* dimensions that follows a Gaussian distribution: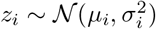, where *µ*_*i*_ and *σ*_*i*_ are the mean and standard deviation of the Gaussian distribution, respectively, and are learned by the encoder network.

The **decoder network p**_*θ*_(**x**|**z**) reconstructs the input features from the latent representations. It includes a fully connected layer, followed by ResNet blocks, and a final convolutional block. The output of this convolutional block is a reconstructed image 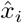 matching the size of the input image.

The **regressor network q**_*ψ*_(**y**|**z**) is designed to predict the dependent variable from the latent representations. It is structured as a simple linear layer that takes in a latent vector *z*_*i*_ and outputs a scalar value *ŷ*_*i*_ representing the predicted dependent variable: *ŷ*_*i*_ = *Wz*_*i*_ + *b*, where *W* and *b* are the learnable weights and bias of the linear layer, respectively.

In the Bayesian framework, the parameters of the encoder (*ϕ*), decoder (*θ*), and regressor (*ψ*) networks are treated as random variables with prior distributions *p*(*ϕ*), *p*(*θ*), and *p*(*ψ*), respectively. Variational inference is employed to approximate the posterior distributions of these parameters given the observed data, by introducing variational distributions *q*(*ϕ*), *q*(*θ*), and *q*(*ψ*). Finally, due to the comparably small training data, instead of learning features from MRI scans directly, in this study we focused on using preprocessed morphometric features as inputs **x** to the VAE (see [32] for processing pipeline. Specifically, we utilized normalized, unmodulated GM, WM, and CSF tissue probability maps. The VAE model learns a m-dimensional latent representation of the sMRI images, which we refer to as the SMAS indices.Given the model construction and the training sample (see next section) we expect the SMAS indices to learn a condensed representation of cognition-related atrophy patterns.

### B. Model optimization basics

In the Bayesian sVAE framework, the primary goal is to optimize a composite loss function that incorporates aspects of Bayesian inference, such as the reconstruction and KL divergence terms from the ELBO, combined with a supervised learning objective. Optimization is conducted using backprop-agation. The loss function is as follows:

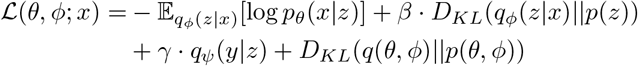

where *θ, ϕ*, and *ψ* denote the parameters of the decoder, encoder, and regressor, respectively. It is composed of three components: reconstruction loss, KL divergence terms, and regression loss. The reconstruction loss, 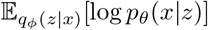, evaluates the sVAE’s ability to accurately reconstruct input data *x* from latent variables *z*, thereby ensuring that the decoded samples bear a close resemblance to the original input data. The KL-divergence terms quantify the disparity between two distributions. The second term, *D*_*KL*_(*q*_*ϕ*_(*z*|*x*)||*p*(*z*)), calculates the divergence between the learned latent variable distribution and a pre-established prior distribution, encouraging the encoder to yield latent variables that align with the prior distribution. The fourth term, *D*_*KL*_(*q*(*θ, ϕ*)||*p*(*θ, ϕ*)), assesses the divergence between the variational distribution of the parameters and their prior distribution. Lastly, the regression loss, *q*_*ψ*_(*y*|*z*), corresponds to the loss associated with the regression task to increase the model’s relevance for cognitive decline. The hyperparameters *β* and *γ* are employed to balance the relative importance of each term in the loss function, with their optimal values generally determined via grid search on out-of-sample data. The selection of appropriate weight priors for the Bayesian sVAE is a challenging task [42]. In our approach, the MOdel Priors with Empirical Bayes using DNN (MOPED) method, as proposed by [42], is applied to establish well-informed weight priors in Bayesian neural networks. To minimize the loss function, the Adam optimization algorithm is utilized [35].

### C. Application to real MRI sample

The DELCODE cohort is a multi-centric observational study conducted at 10 sites of the DZNE. At baseline, the study included 1,079 participants representing a broad spectrum from healthy individuals to those clinically diagnosed with dementia. Specifically, the cohort consisted of 236 CN without any cognitive impairment, 444 subjects with SCD, 191 cases with MCI, 126 AD patients, and 82 ADR (for more details refer [37]). Of the 1,079 participants, 973 subjects (aged 60-89) had T1-weighted MRI scans. Thirty subjects were excluded due to MR artifacts and poor processing quality (see also section A), leaving 943 subjects available for analysis. Longitudinal data included 705 scans at the first annual followup (M12), 550 scans at the second annual follow-up (M24), and scans subjects at the third annual follow-up (M36) (see [13] for more details).

The primary objective of the validation analysis was to evaluate how well the novel SMAS index tracks disease progression and staging, and to validate this index against simpler or alternative markers such as the SPARE-AD index [16]. In addition to testing the model on longitudinal DELCODE data, we further assessed its generalizability using an independent subsample from the ADNI cohort. In the current work, a total of 200 subjects with T1-weighted scans acquired on 1.5 Tesla MRI scanners were retrieved from the ADNI database (as specified by [38]). The sample consisted of 50 subjects with a stable diagnosis of CN state over the 24 months of follow-up, 50 subjects with a stable diagnosis of mild cognitive impairment (sMCI), 50 subjects with a stable diagnosis of AD, and 50 subjects with an initial diagnosis of MCI who showed progression to AD (pMCI) during the follow-up period.

### D. Model training & hyperparameter optimization

We first trained a deterministic sVAE on 75% of the DELCODE baseline sample, conditioned on PACC5. This step derived initial weights for the Bayesian-sVAE, using the MOPED methodology (see above). These pre-trained weights were then used to initialize the Bayesian-sVAE parameters. This approach enhances model convergence and performance by probabilistically modeling uncertainties in both the data and model parameters. The latent representations from the model correspond to SMAS indices. Choosing the dimensionality m of the latent space for VAEs is a challenging task, as it affects all terms in the loss function, e.g., reconstruction error, but particularly interpretability. Our approach focused on optimizing the latent space for tracking subjects on their progression towards AD. Therefore, we decided to identify the number of latent dimensions (SMAS) using a predictive task discriminating between CN and AD subjects, assessed by the area under the receiver operating characteristic curve (AUC-ROC). The performance of the predictive task was evaluated on unseen DELCODE scans at the 12-month follow-up. While a marginal improvement in performance was observed with increased latent dimensions, this difference was not statistically significant, favoring the selection of a single latent dimension for simplicity and interpretability (see supplementary for more details). Finally, a grid search was employed to optimize hyperparameters *β* and *γ*, yielding values of 0.1 and 1, respectively on out-of-sample data. These values were chosen for their ability to maximize predictive performance while maintaining model stability during training. We used the optimal hyperparameters in our further analysis.

### E Model Evaluation

#### 1) Validation of SMAS index: Correlation with Clinical Assessments

To evaluate the effectiveness of the SMAS index in quantifying brain atrophy linked to AD progression, we derive the SMAS index across various validation datasets utilizing the optimal hyperparameters established during the training phase (see section II-D for more details). The SMAS index was normalized with min-max scaling, with the scaling factors generated from the training sample used to the validation datasets for ensuring consistency. We hypothesize that the SMAS index, intended to measure the degree and spatial distribution of brain atrophy, ought to correlate with clinical indicators of disease progression, including the pacc5, hippocampal volume, and total gray matter volume. Additionally, we anticipate a higher association between the SMAS index and age in the DELCODE dataset, which is age-specific, compared to the more heterogeneous ADNI dataset.

#### 2) Tracking Longitudinal Progression of SMAS index

To investigate the usefulness of the SMAS index in tracking AD progression, we conducted a longitudinal analysis utilizing data from the DELCODE and ADNI cohorts. To do this, we applied a linear mixed-effects model to assess temporal changes in the SMAS index across various diagnostic groups (CN, SCD, MCI, & AD).

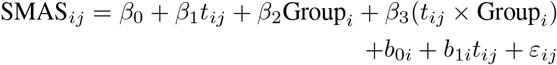

where SMAS_*ij*_ is the SMAS index for subject *i* at timepoint *j, t*_*ij*_ is the time, and Group*i* is the diagnostic group. This aimed to demonstrate whether the SMAS index effectively distinguishes disease stages and captures AD progression dynamics. Additionally, we assessed individual rates of change in the SMAS index and compared them with the rates of atrophy in brain regions typically affected by AD, such as the hippocampus, amygdala, thalamus, ventricles, and total gray matter volume. By comparing these rates, we attempt to demonstrate the association between the SMAS index and region-specific atrophy rates, highlighting the index’s capacity to identify brain atrophy patterns associated with AD progression.

#### 3) Comparative Analysis: SMAS vs. SPARE-AD Index and Hippocampal Volume

To further validate the SMAS index as a marker of brain atrophy, we examined its ability to distinguish between CN, MCI, and AD individuals. We also compared the performance of SMAS index against SPARE-AD index and hippocampal volume in differentiating CN, MCI, and AD in the DELCODE dataset, and CN, stable MCI (sMCI), and progressive MCI (pMCI) in the ADNI dataset. SPARE-AD indices were computed using available code from https://github.com/CBICA/spare_score, while hippocampal volume was obtained using Freesurfer (http://surfer.nmr.mgh.harvard.edu/).

#### 4) Model Transparency and Explainability

To enhance clinical acceptance, we improved SMAS index explainability using gradient activation maps to highlight influential sMRI regions affecting predictions. This approach increases transparency and interpretability, aiming to build trust in model predictions and support informed clinical decision-making.

## III. Results

### A. Validation of SMAS index: Correlating with clinical assessments

We observed significant correlations between SMAS index and various clinical assessments. Higher SMAS scores were associated with cognitive impairment, while lower scores indicated cognitively unimpaired subjects. In the DELCODE dataset (Fig. 2 A), SMAS negatively correlated with PACC5 (r = -0.83), suggesting higher SMAS scores (indicating greater brain atrophy) are associated with lower cognitive performance. Age correlated positively with SMAS (r = 0.50), and correlations with hippocampus volume and total gray matter volume were r = -0.44 and r = -0.42, respectively, aligning with the expected trends. To assess generalizability, we analyzed SMAS index in the ADNI dataset (Fig. 2 B), finding similar trends. The PACC-SMAS correlation was r = -0.62, reaffirming the negative relationship between cognitive performance and SMAS scores. Age showed a weaker correlation (r = 0.28), while correlations with hippocampus volume (r = -0.66) and total gray matter volume (r = -0.47) were consistent.

**Fig. 1.**
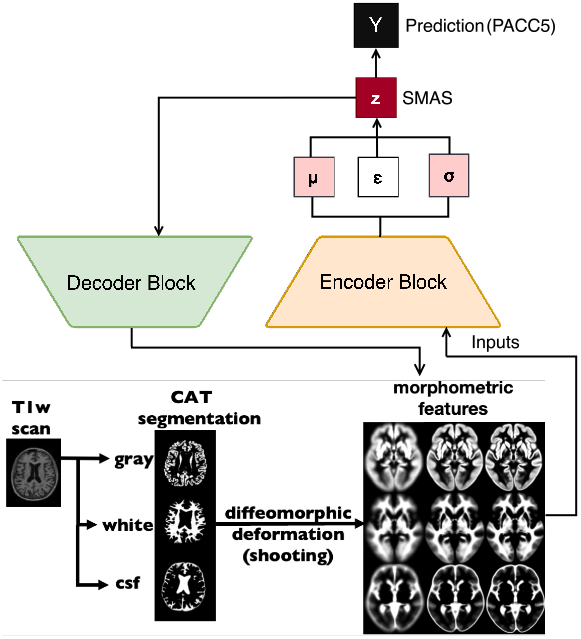
Bayesian-sVAE model for AD-related atrophy patterns: The T1weighted scans are segmented into different brain tissue types (GM, WM & CSF) using the CAT12 segmentation algorithm [32]. The model consists of three main components: an encoder, a prediction block, and a decoder. The encoder encodes the segmented brain tissue features into a latent space to derive the SMAS index. The prediction block uses this latent vector, conditioned on the inputs, to predict cognitive performance differences. Finally, the decoder reconstructs the input space from the latent vector, ensuring the latent representation also captures the original features.

**Fig. 2.**
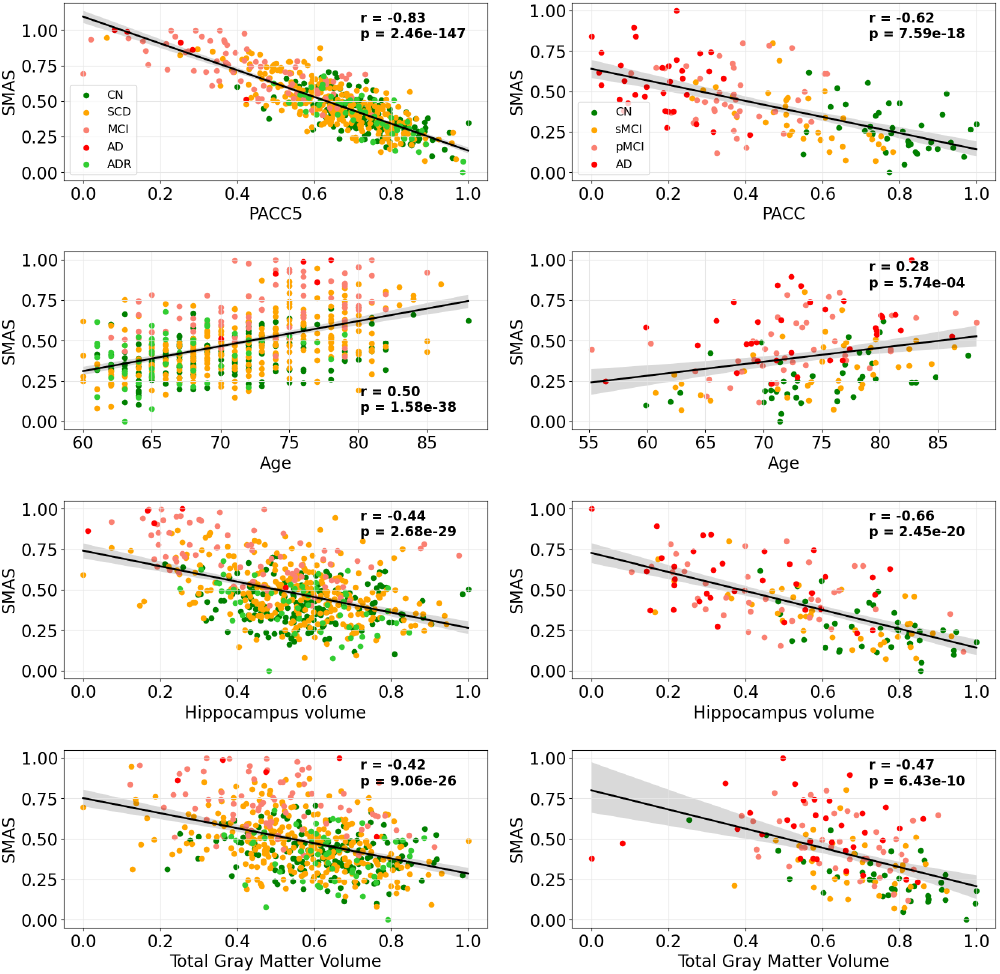
Correlation analysis between SMAS index and clinical measures in (A) DELCODE and (B) ADNI; SMAS shows significant negative correlations with PACC, hippocampus volume, and total gray matter volume, and a positive correlation with age, validating its association with cognitive impairment and brain atrophy.

### B. Longitudinal progression of SMAS

Following the validation of SMAS index, we applied the trained model to analyze the DELCODE and ADNI datasets longitudinally to track disease progression. In the DELCODE dataset (Table I A), includes 1,474 observations across 415 groups, significant differences in disease progression trajectories were observed relative to the CN group. AD subjects exhibited the steepest progression (coefficient = 3.987, p < 0.001), followed by MCI (coefficient = 1.887, p < 0.001), and SCD (coefficient = 0.333, p = 0.002). The progression rates also varied, with AD progressing the fastest (coefficient for time * AD = 0.032, p < 0.001), followed by MCI (0.014, p < 0.001) and SCD (0.007, p < 0.001). Likewise, the ADNI dataset (Table I B), with 759 observations across 190 groups, showed similar patterns. Significant differences in progression were noted compared to the CN reference group, with the AD group showing the highest progression (coefficient = 1.363, p < 0.001), followed by pMCI (0.953, p < 0.001) and sMCI (0.399, p = 0.002). The fastest progression rates were in AD (0.020, p < 0.001), followed by pMCI (0.016, p < 0.001) and sMCI (0.006, p < 0.001).

**TABLE I.**
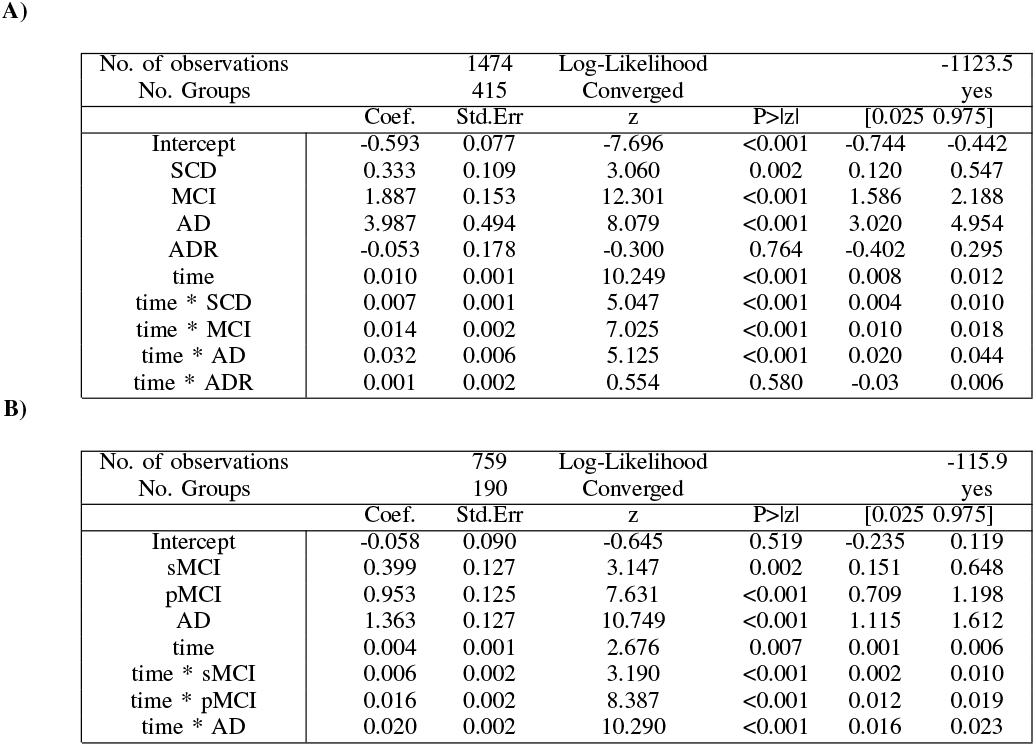
Longitudinal validation of SMAS index across different clinical groups, highlighting the progression trajectories. A) DELCODE dataset B) ADNI dataset. The findings indicate a more rapid progression in AD, succeeded by MCI and SCD individuals.

We also explored the relationship between changes in SMAS index and brain volume changes. In the DELCODE cohort, the rate of change in the SMAS index was significantly and negatively correlated with hippocampal volume change (r = -0.55, p < 0.001), indicating that higher SMAS index corresponds to reduced hippocampal volume. Similar negative & significant correlations were observed for the thalamus (r = -0.32, p < 0.001, p < 0.001), amygdala (r = -0.50, p < 0.001), and total gray matter volume (r = -0.39, p < 0.001), suggesting higher rate of change in SMAS index are linked with higher volume declines in these regions (see supplementary Fig. 3). In the ADNI dataset, a negative correlation between SMAS index change and hippocampal volume change (r = -0.32, p < 0.001) was consistent with the findings from the DELCODE dataset. Additionally, a positive correlation was found with ventricular volume change (r = 0.51, p < 0.001), aligning with typical AD patterns where increased ventricular volume accompanies brain tissue loss (see supplementary Fig. 4). Furthermore, we compared the rate of change in SMAS indices in relation to amyloid beta (A*β*42/40) and phosphorylated tau (p-tau) and observed significant differences between positive and negative groups (see supplementary Fig. 5).

We further examined the longitudinal progression of SMAS in the DELCODE sample across clinical groups and age ranges. The analysis showed that SCD and MCI groups exhibited faster atrophy compared to CN and ADR. When comparing SMAS changes between cognitively unimpaired (CU) and cognitively impaired (CI) groups longitudinally, we observed higher progression rates in the older age group (75-85 years) and in CI individuals across all ages (see supplementary Fig. 2).

### C. Comparative analysis: SMAS vs. SPARE-AD indices and Hippocampus volume

#### 1) DELCODE

We compared the SMAS index with SPARE-AD and hippocampal volume in distinguishing AD, MCI, and CN individuals over time. The SMAS index consistently outperformed across all timepoints.

For CN vs AD (Fig. 3 A), SMAS AUC values remained high: 0.971 at baseline, 0.972 at 12 months, 0.875 at 24 months, and 0.833 at 36 months. SPARE-AD declined from 0.864 at baseline to 0.5 at 24 and 36 months. Hippocampal volume showed a similar drop, from 0.82 to 0.5 by 36 months. In CN vs MCI (Fig. 3 C), SMAS exhibited superior AUC values, from 0.849 at baseline to 0.706 at 36 months. SPARE-AD dropped from 0.751 to 0.605, while hippocampal volume declined from 0.686 to 0.5.

**Fig. 3.**
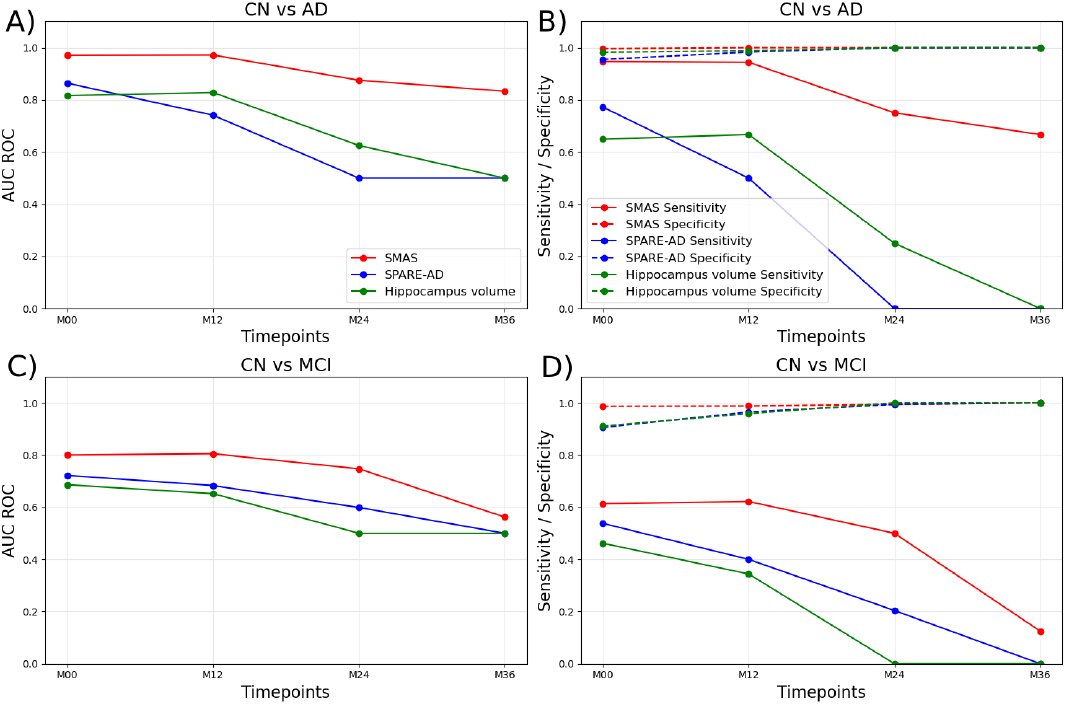
DELCODE: Performance of Hippocampus volume, SPARE-AD, and SMAS index in predicting cognitively normal (CN) vs Alzheimer’s disease (AD) and CN vs mild cognitive impairment (MCI) across different timepoints. (A) Area under the ROC curve (AUC) for CN vs AD prediction. (B) Sensitivity and specificity for CN vs AD prediction. (C) AUC for CN vs MCI prediction. (D) Sensitivity and specificity for CN vs MCI prediction. The SMAS index (red) demonstrates higher AUC, sensitivity and specificity compared to SPARE-AD (blue) and Hippocampus volume (green) for both prediction tasks over the 36-month period

Sensitivity was higher for SMAS in both tasks (Fig. 3). For CN vs AD, it decreased from 0.947 at baseline to 0.667 at 36 months, while SPARE-AD and hippocampal volume dropped to 0 at 24 months. In CN vs MCI, SMAS sensitivity fell from 0.788 to 0.438, outperforming SPARE-AD (0.575 to 0.229) and hippocampal volume (0.462 to 0). Specificity remained high for SMAS, reaching 1.0 for CN vs AD from 12 months onward and improving in CN vs MCI from 0.911 to 0.974 by 36 months. SPARE-AD and hippocampal volume achieved similar specificity after 24 months (Fig. 3 D).

#### 2) ADNI

In the ADNI dataset, SMAS continued to outperform SPARE-AD and hippocampal volume.

For CN vs AD, SMAS AUC values increased from 0.82 at baseline to 0.90 at 24 months, with sensitivity rising from 0.81 to 0.92 and specificity consistently above 0.83. SPARE-AD showed lower AUC (0.66 to 0.75), while hippocampal volume had intermediate values (0.752 to 0.81) (Fig. 4 A, B). For CN vs pMCI, SMAS showed higher AUC (0.76 to 0.812) compared to SPARE-AD (0.69 to 0.68) and hippocampal volume (0.73 to 0.79) (Fig. 4 C, D). In CN vs sMCI, SMAS AUC ranged from 0.625 to 0.62, performing similarly to SPARE-AD and better than hippocampal volume (Fig. 4 E, F).

**Fig. 4.**
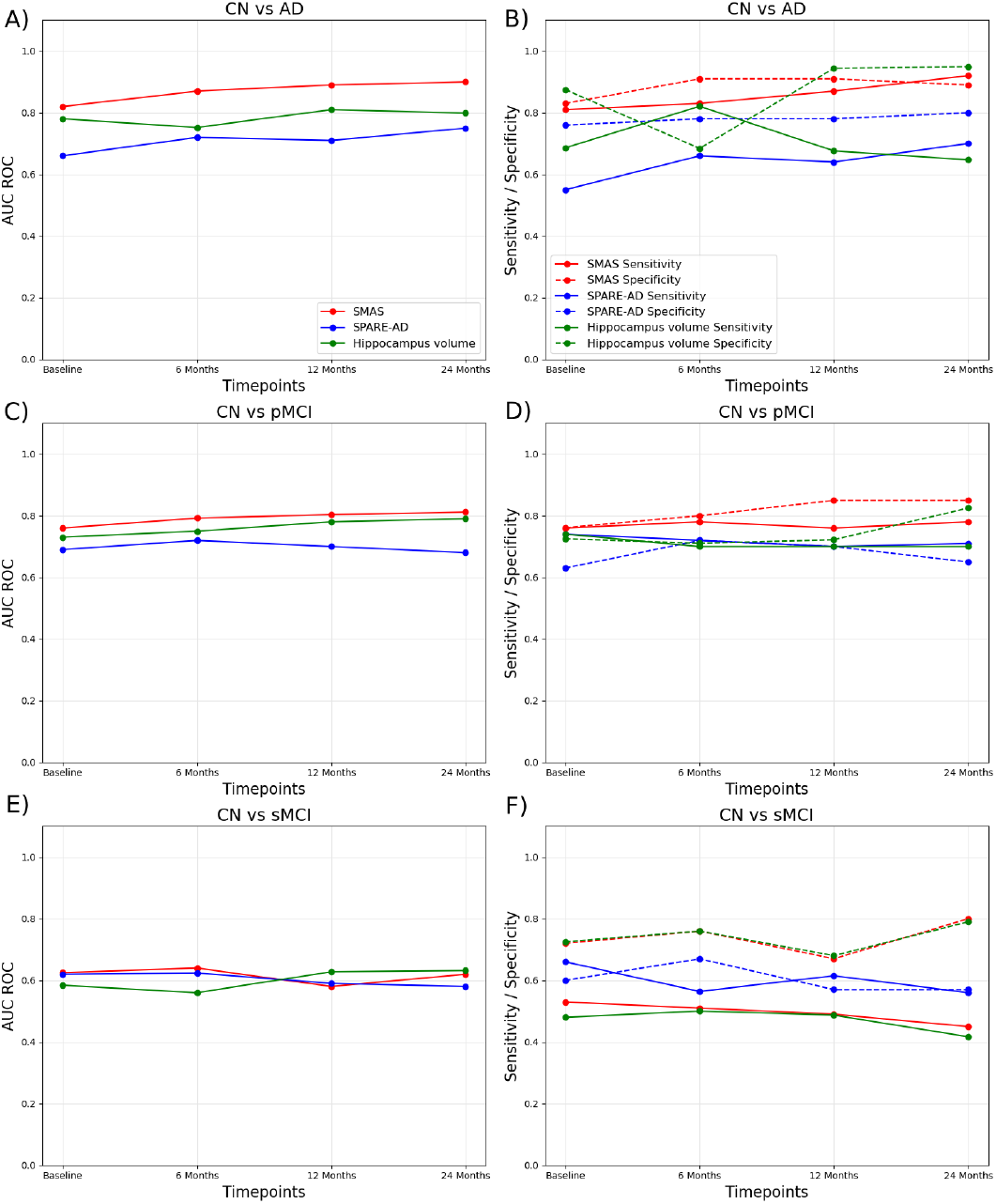
ADNI: Performance of hippocampal volume, SPARE-AD, and SMAS index in predicting CN vs. AD, CN vs. pMCI, and CN vs. sMCI at various time points using an independent ADNI dataset. (A) For CN vs. AD prediction, the SMAS index (red) consistently showed slightly higher performance compared to SPARE-AD (blue) and hippocampal volume (green) over 24 months. (B) Sensitivity and specificity for CN vs. AD prediction further support the favorable performance of the SMAS index. (C) For CN vs. pMCI prediction, the SMAS index exhibited slightly higher AUC values than SPARE-AD and hippocampal volume. (D) Sensitivity and specificity for CN vs. pMCI prediction also suggest a slight advantage for the SMAS index. (E) For CN vs. sMCI prediction, the SMAS index maintained moderately higher ROC AUC values. (F) Sensitivity and specificity for CN vs. sMCI prediction suggest a tendency for higher sensitivity for the SMAS index, while all models demonstrated reasonable specificity over time.

### D. Model transparency

In our study, we examined the relevant maps to derive SMAS index, focusing on the DELCODE M12 validation data. These maps were created by averaging the weight contributions from all subjects in the DELCODE M12 validation dataset, representing the average weight distribution and highlighting regions with varying contributions to the model’s predictions. Our findings reveal the hippocampus, posterior cingulate cortex, precuneus, and lateral parietal cortex as regions with high relevance contributions in estimating SMAS index (Fig. 5). Nevertheless, while these regions demonstrate significant model-based relevance, it is crucial to exercise caution when interpreting these findings. Clinical applicability must be rigorously validated through further empirical research and thorough clinical evaluation.

**Fig. 5.**
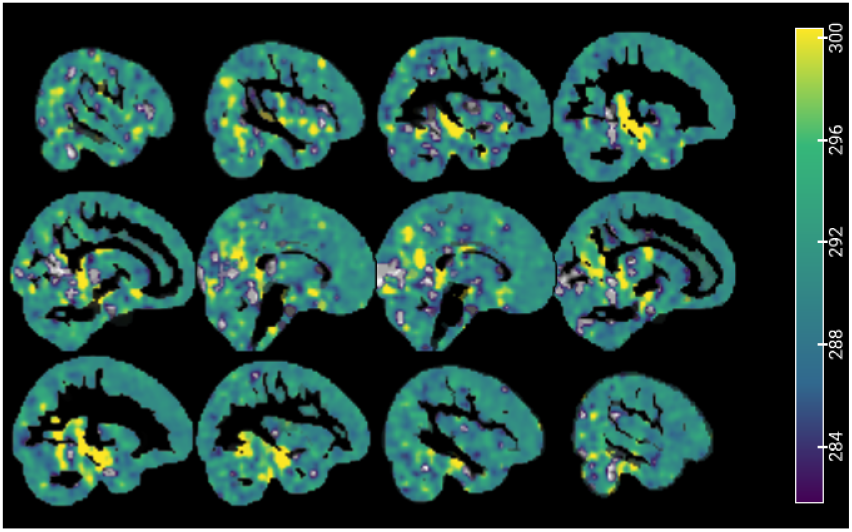
Relevance map of brain regions (hippocampus, posterior cingulate cortex, precuneus, and lateral parietal cortex) contribution in estimating SMAS index

## IV. Discussion

In this study, we present the SMAS index, a novel measure of AD-related morphometric patterns using a deep learning Bayesian-sVAE, trained on structural brain imaging features the large-scale DELCODE cohort. Our results, including a validation of the score across unseen data from two datasets (DELCODE and ADNI), do suggest the robustness and potential of SMAS to characterize at-risk individuals along the progression towards AD.

### A. SMAS index: A reliable indicator of brain atrophy

This study suggests that the SMAS index, derived from a Bayesian-sVAE trained on DELCODE baseline sMRI data, may serve as a useful indicator for assessing brain atrophy. Correlational analyses across independent DELCODE and ADNI datasets reveal associations between the SMAS index and several clinical and structural measures, including PACC5, age, hippocampal volume, and total gray matter volume. A strong negative correlation between SMAS and PACC5 scores suggests that higher SMAS values are associated with lower cognitive performance, indicating potential relevance of SMAS for tracking cognitive decline in AD. Additionally, positive correlations between SMAS and age support its sensitivity to age-related atrophic patterns (see Fig. 2). The SMAS index also demonstrates negative correlations with both hippocampal and total gray matter volumes regions commonly affected in AD indicating its capability to reflect atrophy in these areas [49]–[51]. Longitudinal analyses further support these findings, showing that a higher rate of change in the SMAS index corresponds to reduced volumes in the hippocampus, thalamus, amygdala, and total gray matter, as well as ventricular enlargement. Together, the consistent correlations with cognitive and structural measures, along with the SMAS index sensitivity to longitudinal atrophy changes, highlight its robustness as a reliable indicator of brain atrophy.

### B. SMAS index: A promising tool for early detection and monitoring of AD

#### 1) Monitoring disease progression

The longitudinal analysis of the SMAS index across the DELCODE and ADNI datasets indicates its potential for identifying stages of AD progression. In the DELCODE cohort (n = 1,474 observations over 415 groups), individuals diagnosed with AD demonstrated the highest SMAS progression (*β* = 3.987, p < 0.001) and the fastest rate of change (0.032, p < 0.001). In comparison, those with MCI showed moderate SMAS progression (1.887, p < 0.001) with a rate of change of 0.014 (p < 0.001). SCD participants revealed a more subtle yet significant progression (0.333, p = 0.002) and a slower rate of change (0.007, p < 0.001), confirming the SMAS index sensitivity in identifying early stages of AD. In the ADNI cohort (n = 759 observations across 190 groups), findings were consistent, with AD individuals exhibiting the fastest SMAS progression (1.363, p < 0.001) and the most rapid rate of change (0.020, p < 0.001). pMCI individuals showed higher SMAS progression (0.953, p < 0.001) and a rate of change of 0.016 (p < 0.001), compared to sMCI individuals, who demonstrated lower SMAS progression (0.399, p = 0.002) and a rate of change of 0.006 (p < 0.001). The differences in rates of change between the DELCODE and ADNI cohorts may be attributed to the heterogeneity in the ADNI dataset compared to DELCODE. These results suggest that the SMAS index has potential utility in tracking AD progression.

#### 2) Age-Related Variations

The analysis of the DELCODE cohort reveals varied SMAS trajectories across age and clinical groups, suggesting that SMAS may reflect differences in brain atrophy associated with these groups. In CN individuals, SMAS increases gradually with age, consistent with expected age-related atrophy. In SCD individuals, a steeper SMAS increase is observed, particularly in older age groups. This pattern may suggest that brain atrophy begins to accelerate earlier in SCD, possibly indicating initial neurodegenerative changes before clinical symptoms become apparent [33]. The MCI group shows an even higher SMAS rate from earlier ages, with a steady increase over time, which could suggest that brain atrophy progresses more rapidly in MCI, reflecting a more advanced stage of atrophy. Additionally, CI individuals exhibit a notably steeper SMAS trajectory compared to CU individuals, suggesting an accelerated rate of brain atrophy in CI. Overall, these findings imply that SMAS could be a useful indicator for assessing brain atrophy, with potential utility in identifying and monitoring early stages of AD.

#### 3) Disease Staging

Our analysis of the DELCODE dataset showed that the SMAS index achieved high predictive accuracy in distinguishing CN and AD subjects, demonstrating strong performance with slight decline over time (refer Fig. 3 A). For CN versus MCI, the AUC showed a moderate decrease over time 3 C). The SMAS index outperformed the 3D CNN model by [43], which achieved an AUC of 0.953 for CN vs. AD and 0.775 for CN vs. MCI at baseline.

For the independent ADNI dataset, SMAS index showed strong performance in distinguishing CN from AD and in predicting pMCI to AD conversion, with improving AUC as the time to stable diagnosis decreased. However, the performance for CN vs. sMCI was lower, indicating challenges in differentiating sMCI cases from CN (refer Fig. 3). When compared to other methods, our approach shows competitive performance. For instance, [43] achieved an AUC of 0.949 for AD vs. CN and 0.785 for CN vs. MCI using a 3D CNN. [38], [44], which used a support vector machine with PCA+FDR feature extraction on a similar dataset, reported lower performance metrics, particularly for (CN + sMCI) vs. (pMCI + AD) classification at various time points before stable diagnosis. Additionally, the multi-modal feature selection algorithm (FC2FS) by [46] achieved an AUC of 92.84% for NC vs. AD, while the multi-modal multi-task learning (M3T) method by [47] reported an AUC of 0.933 for AD vs. CN and 0.832 for MCI vs. CN. These comparisons indicate the potential of SMAS index in early disease staging, particularly in distinguishing CN from MCI and pMCI

### C. Comparative analysis

To further validate the robustness of the SMAS index, we compared them with the well-established SPARE-AD and hippocampus volume. We evaluated their ability to differentiate between CN individuals and those with AD or MCI across both the ADNI and DELCODE datasets. Our results consistently showed that the SMAS index demonstrated higher performance compared to SPARE-AD and hippocampus volume over a 36-month observation period. In both datasets, the SMAS model achieved higher AUC scores for CN vs. AD and CN vs. MCI, reaffirming its robustness in early detection (refer Fig. 3, 4).

We also confirmed the effectiveness of the SMAS index against those derived from traditional principal component analysis (PCA). We carried out classification analyses over latent dimensions using an independent subset of the DELCODE dataset (M12). The results echoed our previous findings, showing that the SMAS index consistently outperform traditional methods in distinguishing CN from MCI. This suggests that these index might be useful for the early detection of AD stages.

### D. Transparency and explainability

The quantitative analysis of relevance contributions to SMAS index derived through Bayesian-sVAE demonstrates the model’s capacity to identify and quantify morphological alterations in key brain regions associated with AD. Significant relevance contributions were observed in the hippocampus, posterior cingulate cortex, precuneus, and lateral parietal cortex, aligning with established patterns of AD-related neurodegeneration [60]–[64]. This correlation emphasize the model’s sensitivity to disease-specific structural changes.

Focusing on these critical regions suggests potential for early detection of AD-related atrophy, possibly at the MCI stage; secondly, it offers a means for longitudinal monitoring of disease progression; and thirdly, it presents a promising metric for evaluating the efficacy of potential disease-modifying therapies in clinical trials. This approach offers a transparent and interpretable method for quantifying AD-related brain atrophy.

### E. Limitations and future directions

While our study shows promise, we acknowledge certain limitations and suggest key directions for future work. The current model is primarily based on structural MRI, limiting multimodality to different tissue classes. This choice was based on the increased availability of T1-weighted data within the DELCODE cohort. In future work, we aim to incorporate additional imaging modalities such as fMRI, PET, or DTI, as well as extended demographic and clinical test scores, which could offer a more comprehensive representation of at-risk individuals and capture heterogeneity. This also highlights an important future challenge: addressing the impact of missing data for specific inputs and quantifying uncertainty in latent scores. Another limitation is the use of MRI from multiple imaging sites without explicit post-hoc harmonization to mitigate potential site-specific effects. Although protocols were harmonized a priori in this study, additional variability in MRI acquisition (hardware and software differences) across sites may introduce biases and noise that could affect individual assessments using SMAS in unseen subjects. Future studies might incorporate scan quality indicators and scan-related parameters to enhance the robustness of the score for single subjects. Finally, the potential influence of reserve variables should not be overlooked. Future research may aim to study a wider range of subject-specific factors that could mitigate the effects of brain pathology on cognitive outcomes, especially in assessing an individual’s risk of disease progression.

## References

[1] Nichols, E., Steinmetz, J., Vollset, S., Fukutaki, K., Chalek, J., AbdAllah, F., Abdoli, A., Abualhasan, A., Abu-Gharbieh, E., Akram, T. & Others Estimation of the global prevalence of dementia in 2019 and forecasted prevalence in 2050: an analysis for the Global Burden of Disease Study 2019. The Lancet Public Health. 7, e105–e125 (2022)

[2] Shin, J. Dementia epidemiology fact sheet 2022. Annals Of Rehabilitation Medicine. 46, 53 (2022)

[3] Prince, M., Bryce, R., Albanese, E., Wimo, A., Ribeiro, W. & Ferri, C. The global prevalence of dementia: a systematic review and metaanalysis. Alzheimer’s & Dementia. 9, 63–75 (2013)

[4] Ulep, M., Saraon, S. & McLea, S. Alzheimer disease. The Journal For Nurse Practitioners. 14, 129–135 (2018)

[5] Cummings, J., Morstorf, T. & Zhong, K. Alzheimer’s disease drug-development pipeline: few candidates, frequent failures. Alzheimer’s Research & Therapy. 6 pp. 1-7 (2014)

[6] Blennow, K., Dubois, B., Fagan, A., Lewczuk, P., De Leon, M. & Hampel, H. Clinical utility of cerebrospinal fluid biomarkers in the diagnosis of early Alzheimer’s disease. Alzheimer’s & Dementia. 11, 58–69 (2015)

[7] Qiu, S., Miller, M., Joshi, P., Lee, J., Xue, C., Ni, Y., Wang, Y., De Anda-Duran, I., Hwang, P., Cramer, J. & Others Multimodal deep learning for Alzheimer’s disease dementia assessment. Nature Communications. 13, 3404 (2022)

[8] Jack Jr, C., Bennett, D., Blennow, K., Carrillo, M., Dunn, B., Haeberlein, S., Holtzman, D., Jagust, W., Jessen, F., Karlawish, J. & Others NIA-AA research framework: toward a biological definition of Alzheimer’s disease. Alzheimer’s & Dementia. 14, 535–562 (2018)

[9] Del Sole, A., Malaspina, S. & Biasina, A. Magnetic resonance imaging and positron emission tomography in the diagnosis of neurodegenerative dementias. Functional Neurology. 31, 205 (2016)

[10] Frisoni, G., Fox, N., Jack Jr, C., Scheltens, P. & Thompson, P. The clinical use of structural MRI in Alzheimer disease. Nature Reviews Neurology. 6, 67–77 (2010)

[11] Westman, E., Muehlboeck, J. & Simmons, A. Combining MRI and CSF measures for classification of Alzheimer’s disease and prediction of mild cognitive impairment conversion. Neuroimage. 62, 229–238 (2012)

[12] Jack Jr, C., Petersen, R., Xu, Y., Waring, S., O’Brien, P., Tangalos, E., Smith, G., Ivnik, R. & Kokmen, E. Medial temporal atrophy on MRI in normal aging and very mild Alzheimer’s disease. Neurology. 49, 786–794 (1997)

[13] Nemali, A., Vockert, N., Berron, D., Maas, A., Bernal, J., Yakupov, R., Peters, O., Gref, D., Cosma, N., Preis, L. & Others Gaussian Process-based prediction of memory performance and biomarker status in ageing and Alzheimer’s disease—A systematic model evaluation. Medical Image Analysis. 90 pp. 102913 (2023)

[14] Mateos-Pérez, J., Dadar, M., Lacalle-Aurioles, M., Iturria-Medina, Y., Zeighami, Y. & Evans, A. Structural neuroimaging as clinical predictor: A review of machine learning applications. NeuroImage: Clinical. 20 pp. 506–522 (2018)

[15] Habes, M., Grothe, M., Tunc, B., McMillan, C., Wolk, D. & Davatzikos, C. Disentangling heterogeneity in Alzheimer’s disease and related dementias using data-driven methods. Biological Psychiatry. 88, 70–82 (2020)

[16] Davatzikos, C., Xu, F., An, Y., Fan, Y. & Resnick, S. Longitudinal progression of Alzheimer’s-like patterns of atrophy in normal older adults: the SPARE-AD index. Brain. 132, 2026–2035 (2009)

[17] Ziegler, G., Dahnke, R., Winkler, A. & Gaser, C. Partial least squares correlation of multivariate cognitive abilities and local brain structure in children and adolescents. NeuroImage. 82 pp. 284-294 (2013)

[18] Mihalik, A., Chapman, J., Adams, R., Winter, N., Ferreira, F., Shawe-Taylor, J., Mourão-Miranda, J., Initiative, A. & Others Canonical correlation analysis and partial least squares for identifying brain–behavior associations: A tutorial and a comparative study. Biological Psychiatry: Cognitive Neuroscience And Neuroimaging. 7, 1055–1067 (2022)

[19] Lorenzi, M., Gutman, B., Hibar, D., Altmann, A., Jahanshad, N., Thompson, P. & Ourselin, S. Partial least squares modelling for imaging-genetics in Alzheimer’s disease: Plausibility and generalization. 2016 IEEE 13th International Symposium On Biomedical Imaging (ISBI). pp. 838–841 (2016)

[20] Burzynska, A., Garrett, D., Preuschhof, C., Nagel, I., Li, S., Bäckman, L., Heekeren, H. & Lindenberger, U. A scaffold for efficiency in the human brain. Journal Of Neuroscience. 33, 17150–17159 (2013)

[21] Nakua, H., Yu, J., Abdi, H., Hawco, C., Voineskos, A., Hill, S., Lai, M., Wheeler, A., McIntosh, A. & Ameis, S. Comparing the stability and reproducibility of brain-behaviour relationships found using Canonical Correlation Analysis and Partial Least Squares within the ABCD Sample. Network Neuroscience. pp. 1–52 (2024)

[22] Diogo, V., Ferreira, H., Prata, D. & Initiative, A. Early diagnosis of Alzheimer’s disease using machine learning: a multi-diagnostic, generalizable approach. Alzheimer’s Research & Therapy. 14, 107 (2022)

[23] Wang, Y., Gao, R., Wei, T., Johnston, L., Yuan, X., Zhang, Y., Yu, Z. & Initiative, A. Predicting long-term progression of Alzheimer’s disease using a multimodal deep learning model incorporating interaction effects. Journal Of Translational Medicine. 22, 265 (2024)

[24] Venugopalan, J., Tong, L., Hassanzadeh, H. & Wang, M. Multimodal deep learning models for early detection of Alzheimer’s disease stage. Scientific Reports. 11, 3254 (2021)

[25] Ebrahimighahnavieh, M., Luo, S. & Chiong, R. Deep learning to detect Alzheimer’s disease from neuroimaging: A systematic literature review. Computer Methods And Programs In Biomedicine. 187 pp. 105242 (2020)

[26] Jo, T., Nho, K. & Saykin, A. Deep learning in Alzheimer’s disease: diagnostic classification and prognostic prediction using neuroimaging data. Frontiers In Aging Neuroscience. 11 pp. 220 (2019)

[27] Kingma, D., Welling, M. & Others An introduction to variational autoencoders. Foundations And Trends® In Machine Learning. 12, 307–392 (2019)

[28] Kingma, D., Mohamed, S., Jimenez Rezende, D. & Welling, M. Semi-supervised learning with deep generative models. Advances In Neural Information Processing Systems. 27 (2014)

[29] Sauty, B. & Durrleman, S. Progression models for imaging data with longitudinal variational auto encoders. International Conference On Medical Image Computing And Computer-Assisted Intervention. pp. 3–13 (2022)

[30] Basu, S., Wagstyl, K., Zandifar, A., Collins, L., Romero, A. & Precup, D. Early prediction of alzheimer’s disease progression using variational autoencoders. Medical Image Computing And Computer Assisted In-tervention–MICCAI 2019: 22nd International Conference, Shenzhen, China, October 13–17, 2019, Proceedings, Part IV 22. pp. 205–213 (2019)

[31] Zhao, Q., Adeli, E., Honnorat, N., Leng, T. & Pohl, K. Variational autoencoder for regression: Application to brain aging analysis. Medical Image Computing And Computer Assisted Intervention–MICCAI 2019: 22nd International Conference, Shenzhen, China, October 13–17, 2019, Proceedings, Part II 22. pp. 823-831 (2019)

[32] Gaser, Christian, Robert Dahnke, Paul M. Thompson, Florian Kurth, Eileen Luders, and Alzheimer’s Disease Neuroimaging Initiative. CAT: a computational anatomy toolbox for the analysis of structural MRI data. GigaScience 13 (2024): giae049.

[33] Lerch, O., Ferreira, D., Stomrud, E., van Westen, D., Tideman, P., Palmqvist, S., Mattsson-Carlgren, N., Hort, J., Hansson, O. and Westman, E., 2024. Predicting progression from subjective cognitive decline to mild cognitive impairment or dementia based on brain atrophy patterns. Alzheimer’s Research Therapy, 16(1), p.153.

[34] Huang, L., Ruan, S., Xing, Y. & Feng, M. A review of uncertainty quantification in medical image analysis: probabilistic and non-probabilistic methods. (2023)

[35] Jais, I.K.M., Ismail, A.R. and Nisa, S.Q., 2019. Adam optimization algorithm for wide and deep neural network. Knowl. Eng. Data Sci., 2(1), pp.41–46.

[36] Lambert, B., Forbes, F., Tucholka, A., Doyle, S., Dehaene, H. & Dojat, M. Trustworthy clinical AI solutions: a unified review of uncertainty quantification in deep learning models for medical image analysis. (2022)

[37] Jessen, F., Spottke, A., Boecker, H., Brosseron, F., Buerger, K., Catak, C., Fliessbach, K., Franke, C., Fuentes, M., Heneka, M. & Others Design and first baseline data of the DZNE multicenter observational study on predementia Alzheimer’s disease (DELCODE). Alzheimer’s Research & Therapy. 10 pp. 1–10 (2018)

[38] Salvatore, C., Cerasa, A. & Castiglioni, I. MRI characterizes the progressive course of AD and predicts conversion to Alzheimer’s dementia 24 months before probable diagnosis. Frontiers In Aging Neuroscience. 10 pp. 135 (2018)

[39] Tohka, J., Zijdenbos, A. & Evans, A. Fast and robust parameter estimation for statistical partial volume models in brain MRI. Neuroimage. 23, 84–97 (2004)

[40] Gaser, C. Partial volume segmentation with adaptive maximum a posteriori (MAP) approach. NeuroImage. 47 pp. S121 (2009)

[41] Ashburner, J. & Friston, K. Diffeomorphic registration using geodesic shooting and Gauss–Newton optimisation. NeuroImage. 55, 954–967 (2011)

[42] Krishnan, R., Subedar, M. & Tickoo, O. Specifying Weight Priors in Bayesian Deep Neural Networks with Empirical Bayes. (2019)

[43] Dyrba, M., Hanzig, M., Altenstein, S., Bader, S., Ballarini, T., Brosseron, F., Buerger, K., Cantré, D., Dechent, P., Dobisch, L. & Others Improving 3D convolutional neural network comprehensibility via interactive visualization of relevance maps: evaluation in Alzheimer’s disease. Alzheimer’s Research & Therapy. 13 pp. 1–18 (2021)

[44] Salvatore, C., Cerasa, A., Battista, P., Gilardi, M., Quattrone, A., Castiglioni, I. & Initiative, A. Magnetic resonance imaging biomarkers for the early diagnosis of Alzheimer’s disease: a machine learning approach. Frontiers In Neuroscience. 9 pp. 307 (2015)

[45] Jack Jr, Clifford R., David S. Knopman, William J. Jagust, Ronald C. Petersen, Michael W. Weiner, Paul S. Aisen, Leslie M. Shaw et al. Update on hypothetical model of Alzheimer’s disease biomarkers. Lancet neurology 12, no. 2 (2013): 207.

[46] Jiao, Z., Chen, S., Shi, H. & Xu, J. Multi-modal feature selection with feature correlation and feature structure fusion for MCI and AD classification. Brain Sciences. 12, 80 (2022)

[47] Zhang, D., Shen, D., Initiative, A. & Others Multi-modal multi-task learning for joint prediction of multiple regression and classification variables in Alzheimer’s disease. NeuroImage. 59, 895–907 (2012)

[48] He, K., Zhang, X., Ren, S. & Sun, J. Deep residual learning for image recognition. Proceedings Of The IEEE Conference On Computer Vision And Pattern Recognition. pp. 770–778 (2016)

[49] Rao, Y., Ganaraja, B., Murlimanju, B., Joy, T., Krishnamurthy, A. & Agrawal, A. Hippocampus and its involvement in Alzheimer’s disease: a review. 3 Biotech. 12, 55 (2022)

[50] Cvetković-Dožić, D., Skender-Gazibara, M. & Dožić, S. Neuropathological hallmarks of Alzheimer’s disease. Archive Of Oncology. 9, 195–199 (2001)

[51] Mueller, S., Schuff, N., Yaffe, K., Madison, C., Miller, B. & Weiner, M. Hippocampal atrophy patterns in mild cognitive impairment and Alzheimer’s disease. Human Brain Mapping. 31, 1339–1347 (2010)

[52] Stern, Y. What is cognitive reserve? Theory and research application of the reserve concept. Journal Of The International Neuropsychological Society. 8, 448–460 (2002)

[53] Pini, L., Pievani, M., Bocchetta, M., Altomare, D., Bosco, P., Cavedo, E., Galluzzi, S., Marizzoni, M. & Frisoni, G. Brain atrophy in Alzheimer’s disease and aging. Ageing Research Reviews. 30 pp. 25–48 (2016)

[54] Zhou, M., Zhang, F., Zhao, L., Qian, J. & Dong, C. Entorhinal cortex: a good biomarker of mild cognitive impairment and mild Alzheimer’s disease. Reviews In The Neurosciences. 27, 185–195 (2016)

[55] Casanova, R., Barnard, R., Gaussoin, S., Saldana, S., Hayden, K., Manson, J., Wallace, R., Rapp, S., Resnick, S., Espeland, M. & Others Using high-dimensional machine learning methods to estimate an anatomical risk factor for Alzheimer’s disease across imaging databases. Neuroimage. 183 pp. 401–411 (2018)

[56] Bastos-Leite, A., Van Der Flier, W., Van Straaten, E., Staekenborg, S., Scheltens, P. & Barkhof, F. The contribution of medial temporal lobe atrophy and vascular pathology to cognitive impairment in vascular dementia. Stroke. 38, 3182–3185 (2007)

[57] Chan, D., Fox, N., Scahill, R., Crum, W., Whitwell, J., Leschziner, G., Rossor, A., Stevens, J., Cipolotti, L. & Rossor, M. Patterns of temporal lobe atrophy in semantic dementia and Alzheimer’s disease. Annals Of Neurology. 49, 433–442 (2001)

[58] Van De Pol, L., Hensel, A., Flier, W., Visser, P., Pijnenburg, Y., Barkhof, F., Gertz, H. & Scheltens, P. Hippocampal atrophy on MRI in frontotemporal lobar degeneration and Alzheimer’s disease. Journal Of Neurology, Neurosurgery & Psychiatry. 77, 439–442 (2006)

[59] Selvaraju, R., Cogswell, M., Das, A., Vedantam, R., Parikh, D. & Batra, D. Grad-cam: Visual explanations from deep networks via gradient-based localization. Proceedings Of The IEEE International Conference On Computer Vision. pp. 618–626 (2017)

[60] Poulakis, K., Pereira, J., Muehlboeck, J., Wahlund, L., Smedby, Ö., Volpe, G., Masters, C., Ames, D., Niimi, Y., Iwatsubo, T. & Others Multi-cohort and longitudinal Bayesian clustering study of stage and subtype in Alzheimer’s disease. Nature Communications. 13, 4566 (2022)

[61] Yang, H., Xu, H., Li, Q., Jin, Y., Jiang, W., Wang, J., Wu, Y., Li, W., Yang, C., Li, X. & Others Study of brain morphology change in Alzheimer’s disease and amnestic mild cognitive impairment compared with normal controls. General Psychiatry. 32 (2019)

[62] Levin, F., Ferreira, D., Lange, C., Dyrba, M., Westman, E., Buchert, R., Teipel, S., Grothe, M. & Initiative, A. Data-driven FDG-PET subtypes of Alzheimer’s disease-related neurodegeneration. Alzheimer’s Research & Therapy. 13 pp. 1–14 (2021)

[63] Lee, P., Chou, K., Chung, C., Lai, T., Zhou, J., Wang, P. & Lin, C. Posterior cingulate cortex network predicts Alzheimer’s disease progression. Frontiers In Aging Neuroscience. 12 pp. 608667 (2020)

[64] Rami, L., Sala-Llonch, R., Solé-Padullés, C., Fortea, J., Olives, J., Lladó, A., Pena-Gómez, C., Balasa, M., Bosch, B., Antonell, A. & Others Distinct functional activity of the precuneus and posterior cingulate cortex during encoding in the preclinical stage of Alzheimer’s disease. Journal Of Alzheimer’s Disease. 31, 517–526 (2012)

