## Supplementary material for "SMAS: Structural MRI-Based AD Score using Bayesian VAE"

#### 1 DELCODE MRI acquisition

MRI scans were acquired in 9 out of 10 involved DZNE sites (3T Siemens scanners: 3 TIM Trio systems, 4 Verio systems, 1 Skyra and 1 Prisma system). Our main analyses were based on whole-brain T1-weighted MPRAGE (3D GRAPPA PAT 2, 1 mm3 isotropic, 256 X 256 px, 192 slices, sagittal, 5 min, TR 2500 ms, TE 4.33 ms, TI 110 ms, FA 7°). Further ROI and covariate processing was based on the additionally available FLAIR protocol (for details see Jessen, F., et al. 2018).

#### 2 Demographics information

| **Variable** | **CN** | **SCD** | **MCI** | **AD** | **ADR** |
|---|---|---|---|---|---|
| **No. of subjects** | 229 | 388 | 158 | 109 | 75 |
| **Males/females** | 129/94 | 173/200 | 69/82 | 63/44 | 44/31 |
| **Age (Mean ± SD)** | 69.46 ± 5.42 | 71.27 ± 6.06 | 72.91 ± 5.72 | 75.19 ± 6.25 | 66.27 ± 4.61 |
| **PACC5 (Mean ± SD)** | 0.77 ± 0.09 | 0.72 ± 0.11 | 0.49 ± 0.14 | 0.25 ± 0.10 | 0.76 ± 0.11 |

Table 1: Baseline demographic information for the participants from the DELCODE cohort used in this modelling study. The PACC5 score is transformed using min-max normalization to the unit interval. Age is indicated in years.

| **Variable** | **CN** | **sMCI** | **pMCI** | **AD** |
|---|---|---|---|---|
| **No. of subjects** | 50 | 50 | 50 | 50 |
| **Age (Mean ± SD)** | 74.84 ± 6.10 | 74.66 ± 6.83 | 74.49 ± 7.08 | 75.14 ± 6.75 |
| **PACC (Mean ± SD)** | 0.77 ± 0.11 | 0.52 ± 0.12 | 0.39 ± 0.12 | 0.21 ± 0.10 |

Table 2: Baseline demographic information for the participants from the ADNI cohort used in this modelling study. The PACC score is transformed using min-max normalization to the unit interval. Age is indicated in years.

### 3 Results

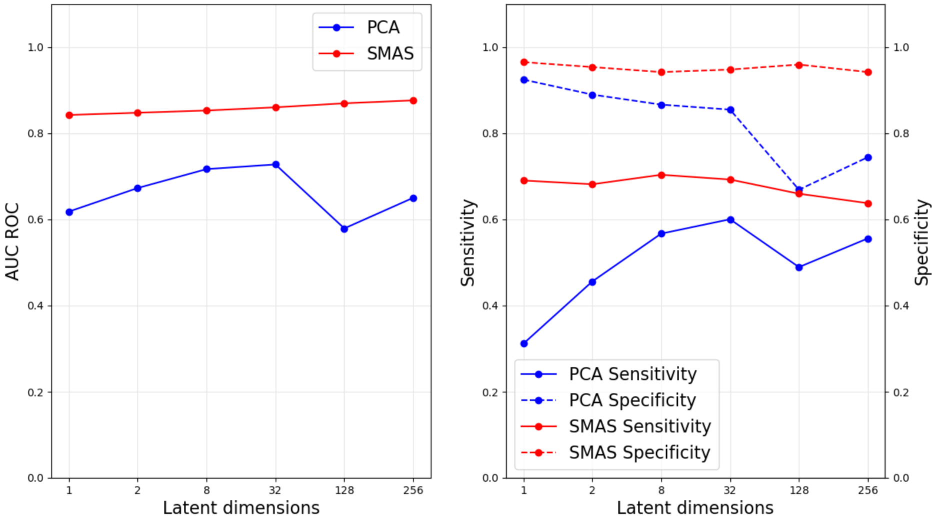


Figure 1: Comparative evaluation of CN vs. MCI classification performance using indices derived from PCA and Bayesian-VAE for the DELCODE M12 sample trained on the DELCODE baseline sample: SMAS indices exhibit superior AUC performance over PCA indices across latent dimensions, demonstrating high specificity and sensitivity. Despite a slight performance enhancement with increasing latent dimensions, the difference was not statistically significant, advocating for the choice of a single latent dimension for ease of interpretation and simplicity.

#### 3.1 Longitudinal rate of change in SMAS indices

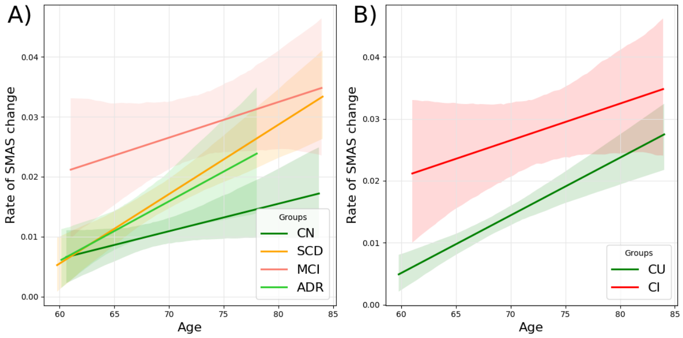


Figure 2: Rate of SMAS change over Age for different clinical groups A) Comparison of the rate of SMAS change across age for CN, SCD, MCI, AD and ADR groups. B) Rate of SMAS change over age for cognitively unimpaired (CU) and cognitively impaired (CI) groups.

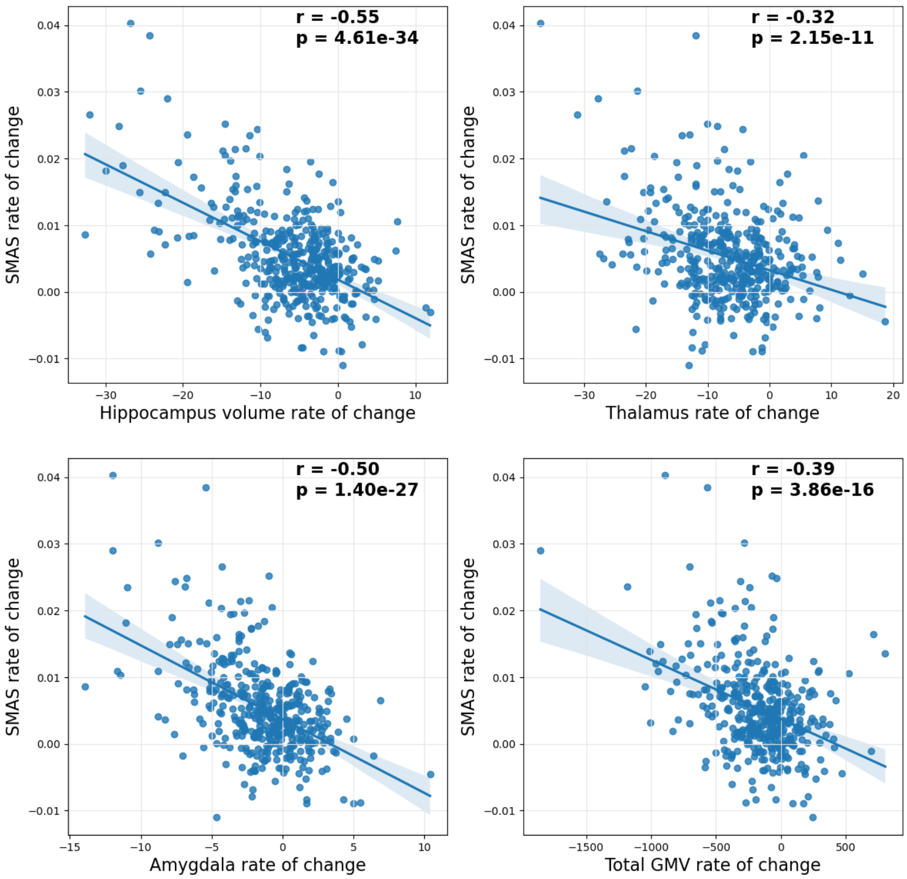


Figure 3: Rate of change of SMAS indices and brain volume changes in DELCODE Sample. The scatter plots depict the correlation between the rate of change of SMAS indices and the rate of change in the volume of the Hippocampus (r = -0.55), Thalamus (r = -0.32), Amygdala (r = -0.50), and total Gray Matter Volume (r = -0.39).

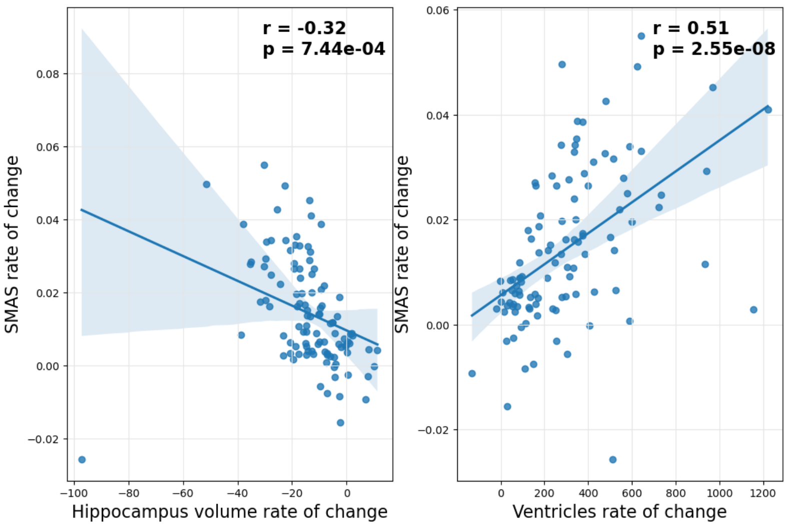


Figure 4: Rate of change of SMAS indices and brain volume changes in ADNI Sample. The scatter plots depict the correlation between the rate of change of SMAS indices and the rate of change in the volume of the Hippocampus (r = -0.32), Ventricles (r = 0.51).

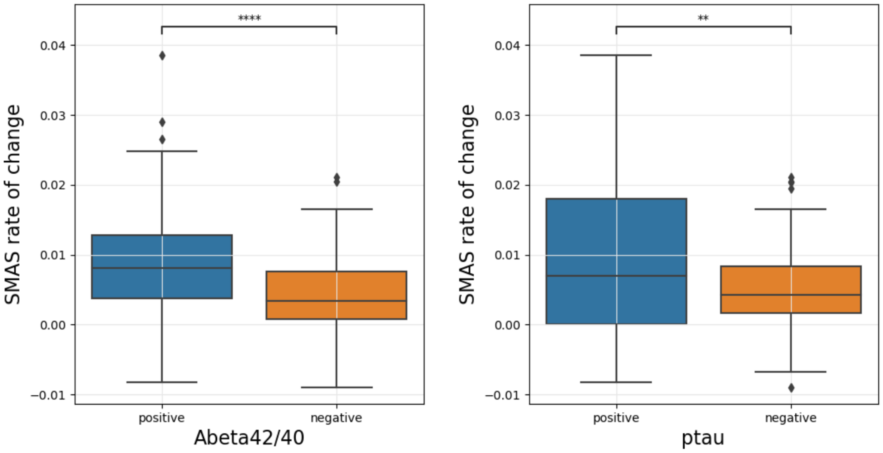


Figure 5: Comparison of SMAS indices rate of change in relation to CSF biomarkers (A$\beta$ 42/40) and ptau. The left side represents the significant rate of change in SMAS indices for both A$\beta$42/40 positive and negative groups. On the right side, for both ptau positive and negative groups, we observe a significant rate of change

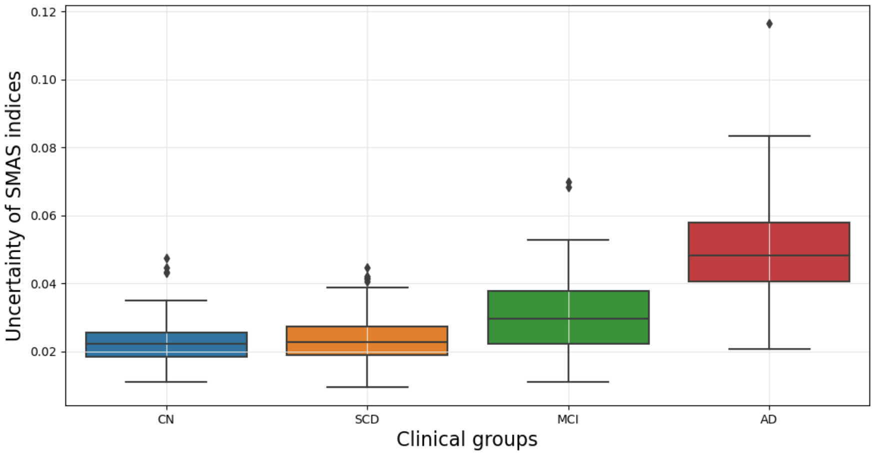


Figure 6: Uncertainty of SMAS indices for DELCODE cohort across clinical groups. The CN and SCD groups exhibit relatively low uncertainty, indicating more consistent SMAS index measurements in these groups. The MCI group shows moderate uncertainty with a broader distribution, reflecting increased variability. The AD group presents the highest level of uncertainty, suggesting significant variability in SMAS indices within this population.

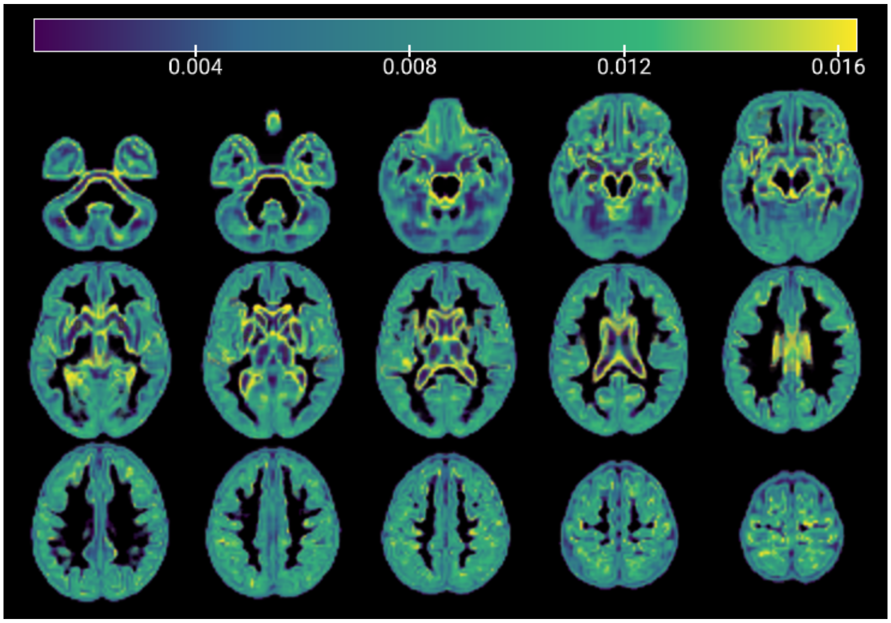


Figure 7: Reconstruction uncertainty from the Bayesian-sVAE model in estimating SMAS indices. Higher uncertainty areas are observed primarily in the central regions around the ventricles, the boundaries and edges of brain structures, and some cortical regions in the lower slices. These areas reflect the regions where the Bayesian-sVAE model has more difficulty in accurate reconstruction, potentially due to the variability in the sMRI.
